# Persistent Behavioral and Physiological Effects of Dietary Protein Restriction

**DOI:** 10.1101/2024.03.04.583396

**Authors:** Paul L. Soto, Christopher D. Morrison

## Abstract

Protein provides essential amino acids critical for survival. Recent research has identified that dietary protein restriction induces physiological and behavioral adaptations and that those adaptations are mediated by liver-produced fibroblast growth factor 21 (FGF21) that acts in the brain. Most of the research on adaptations to dietary protein restriction and the biological factors that mediate those adaptations carries an implicit assumption that the effects of dietary protein restriction are reversible. Rarely is the assumption of reversibility directly examined by varying dietary protein content within animals. Recently collected data on preference for protein versus carbohydrate solutions indicates that when mice are placed on a low (5%) protein diet, preference for protein is rapidly increased. However, when mice are switched from a low to normal (20%) protein diet, relative consumption of protein decreases but preference for protein persists. Re-analysis of published data suggests, similarly, that preference for protein-rich food sources remains elevated following exposure to dietary protein restriction and that calcium influx in the ventral tegmental area of the brain in response to protein consumption also remains persistently elevated following dietary protein restriction. Together, the data suggest that prior protein restriction can exert long-lasting effects on protein preference and on brain activity associated with protein consumption. The finding that the effects of protein restriction do not rapidly reverse implies that protein choice may be a function, not only of current nutritional state, but of historical nutritional conditions. Further, these results suggest that a full understanding of the environmental and biological determinants of protein choice requires studies in which dietary protein intake is varied within subjects.

---

Protein is critical for survival. For rodents, a balanced diet contains around 15-20% protein, by weight. Moderate dietary protein restriction to around 5-10%, by weight, induces physiological and behavioral changes including decreased body weight, increased energy expenditure, increased expression of *Fgf*21 and increased circulating FGF21, increased consumption of low protein food, and increased preference for and consumption of protein-rich food sources (Chaumontet et al., 2019; Chiacchierini et al., 2021; Hill et al., 2017; Hill et al., 2019; Hill et al., 2020; Laeger et al., 2014; Murphy et al., 2018; Naneix et al., 2020; Torres et al., 2022; Volcko & McCutcheon, 2022; White et al., 2000a, 2000b; Zapata et al., 2019). Robust behavioral and physiological adaptations to dietary protein restriction are consistent with the idea of protein leverage, in which animals prioritize protein intake and will overeat other macronutrients to reach a protein target (Raubenheimer & Simpson, 2019; Simpson & Raubenheimer, 2005).

Many studies on the effects of dietary protein restriction involve comparisons of groups of animals fed diets differing in protein content; typically, a “normal” protein diet containing around 14-20% protein and a “low” protein diet containing around 5% protein. Such studies, at a minimum, define how animals differ when maintained on diets of varying protein content. However, such comparisons do not necessarily provide information related to how individual animals respond when protein content is dynamically varied over time, and prior nutritional experiences may affect subsequent responses to nutritional variation.

Studies on the effects of reducing dietary protein content are often interpreted as providing information on how animals dynamically adapt to fluctuations in protein intake. For example, a recent paper on the effect of dietary protein restriction on stimulated dopamine release begins “The regulation of food intake *in an ever-changing environment* [emphasis added] is a central survival process” (Naneix et al., 2021, p. 394). Similarly, a recent study on the effects of dietary protein restriction on food choice and metabolism and the role of FGF21 in those effects begins “The survival of any species is contingent on the ability to physiologically and behaviorally adapt to a *changing nutritional environment* [emphasis added]” (Hill et al., 2019, p. 2934). Our understanding of how animals adapt to changes in dietary protein availability primarily comes from comparisons between groups of animals fed diets of differing protein content. Careful examination of the existing literature and re-analysis of our own data suggest that dietary protein restriction, at least in the moderate range, induces persistent behavioral (and possibly physiological) effects. Thus, it may be that current physiological and behavioral responses related to protein are jointly determined by prior protein restriction and current dietary conditions.

We initially became interested in the issue of within-animal variations in dietary protein intake because of the potential utility of varying dietary protein content within animals (for a discussion of this topic, see Soto, 2020), which is particularly important when behavior requires extended training or preparation of animals requires extensive time and labor (e.g., surgical preparation of animals for fiber photometry studies). In our own attempts to use animals more efficiently, we found, to our surprise, that although behavior adjusted rapidly when animals were initially protein-restricted, changes were slower and occurred to a lesser extent when protein content was normalized following a prior period of protein restriction. Subsequent careful examination of the literature, which we review here, reveals other examples of this phenomenon, which as we have argued above is important for our understanding of the dynamics of behavioral and physiological adaptations to variation in dietary protein content.

## Methods

### Studies Included

Although many studies have compared solid food intake across groups of animals consuming diets of varying protein content, we have only identified one study (Dibattista & Holder, 1998) that allows a comparison of changes in food intake when dietary protein content was normalized in animals that previously experienced dietary protein restriction. DiBattista and Holder (1998) conducted an experiment (Experiment 2) in which rats were fed a protein-free (protein-restricted; PR) or nutritionally complete (not-restricted; NR) maintenance diet for 10 days (22 h/day access), subsequently placed on a nutritionally complete chow diet (Purina Rodent Chow 5001) for 25 days, and then fed either the protein-free or nutritionally complete maintenance diet for 10 more days (22 h/day access). During the first and second 10-day periods, the rats were given daily 2-h test sessions in which they could consume either of two test diets – a protein-rich test diet or a carbohydrate-rich test diet. Four groups of rats were studied: 1) rats fed the protein-free maintenance diet before and after the intervening 25-day period on chow (the PR-PR group), 2) rats that were initially given the protein-free maintenance diet and were given the nutritionally complete maintenance diet after the intervening 25-day period on chow (the PR-NR group), 3) rats that were initially given the nutritionally complete maintenance diet and subsequently the protein-free maintenance diet after the intervening 25-day period on chow (the NR-PR group), and 4) rats initially given the nutritionally complete maintenance diet before and after the intervening 25-day period on chow (the NR-NR group).

We identified three studies in which preference for a protein vs. carbohydrate solution was evaluated during exposure to a normal protein diet and during exposure to a low protein diet in the same animals (Chiacchierini et al., 2021; Naneix et al., 2020; Torres et al., 2022). In the Torres et al. (2022) study, one group of mice was exposed to 21 days with a normal protein (18% casein, by weight) diet (D11051801, Research Diets Inc, New Brunswick, NJ) followed by 28 days with a low protein (4% casein, by weight) diet (D11092301, Research Diets Inc). A second group of mice received 21 days exposure to the low protein diet followed by 28 days exposure to the normal protein diet. In each group, preference for a 4% casein versus 4% maltodextrin was assessed twice during each diet exposure using a 24 h/day, 3-day, two-bottle choice procedure. In the other two preference studies (Chiacchierini et al., 2021; Naneix et al., 2020), one group of rats received 18 days of exposure a normal protein diet (14% casein, by weight; EU Rodent Diet 5LF2, Lab Diet) followed by 14 days of exposure to a low protein diet (5% casein, by weight; D15100602, Research Diets Inc). A second group of rats in each study received 18 days of exposure to the low protein diet followed by 14 days exposure to the normal protein diet. Preference for casein versus maltodextrin solution was assessed on day 18 of exposure to the first diet and at days 7 and 14 of exposure to the second diet using a discrete trials procedure in which licks on each of two concurrently available sipper tubes (casein versus maltodextrin) were counted via computer interface (diet and timing details confirmed via personal communication with James McCutcheon; conditioning sessions with only a single solution were conducted prior to the first and third preference assessments).

In addition to measuring licks on the casein and maltodextrin sipper tubes, Chiacchierini et al. (2021) also used fiber photometry to measure calcium influx in the ventral tegmental area (VTA) when licks occurred on the casein and maltodextrin sipper tubes during forced trials in which only a single sipper tube was provided. Thus, the Chiacchierini et al. (2021) study also provides a measure of VTA activity, quantified as area-under-the-curve (AUC) values in the photometry signal, in response to casein versus maltodextrin under conditions of low and normal protein diet feeding.

### Data Extraction/Collection

Data from Dibattista and Holder (1998) and Naneix et al. (2020) were extracted using the free PlotDigitizer web application (*PlotDigitizer*). Previously published data from our laboratory (Torres et al., 2022, Cohort 2) were re-analyzed and re-plotted to show amounts, in grams, of protein (casein) and carbohydrate (maltodextrin) solution consumed during each of four preference assessments. Data from Chiacchierini et al. (2021) were downloaded from the corresponding author’s publicly available repository (McCutcheon, 2021) and converted into csv files using the Python programming language.

### Data Analysis

Data from DiBattista and Holder (1998) and Naneix et al. (2020) were not analyzed statistically because individual animal data cannot be extracted from the relevant figures. Data from Torres et al. (2022) were analyzed using a linear multilevel model in which amount of solution consumed, in grams, was the dependent variable and Solution (casein or maltodextrin; categorical, dummy-coded with casein as reference level), Diet (Low Protein or Normal Protein; categorical, dummy-coded with Normal Protein as the reference level), Group (Low Protein First or Normal Protein First; categorical, dummy-coded with Normal Protein First as the reference level), and Assessment number (1 or 2 within each diet; continuous) were used as predictors with all possible interactions included. Solution and Diet were also included as random effects at the individual mouse level.

Similarly, data from Chiacchierini et al. (2022) were analyzed using a linear multilevel model with licks or AUC value as the dependent variable and predictors of Group (Low Protein First or Normal Protein First; categorical, dummy-coded with Low Protein First as the reference level), Day (0, 7, or 14; continuous), Solution (casein or maltodextrin; categorical, dummy-coded with casein as the reference level). Only the intercept was included as a random effects predictor at the individual animal level because including other predictors as random effects prevented model convergence. Diet was not included as a predictor because there was only one assessment during the first diet exposure and thus the Day predictor fully encompassed the diet sequence.

All calculations were conducted in the R programming language (R Core Team, 2018), linear multilevel models were conducted using the lmer command in the lme4 package (Bates et al., 2015), and figures were created using the ggplot2 package (Wickham, 2016). The emmeans command of the emmeans package (Lenth, 2020) was used to calculate fixed-effect level marginal means and to assess differences between Groups and Solutions for statistical significance using t-tests with degrees of freedom calculated using the Kenward-Roger method. Multilevel models were selected for analysis because they take into account the nested (repeated measures within each animal) and crossed (observations on animals within multiple conditions) nature of data such as those reported here and provide acceptable type I error rates (Boisgontier & Cheval, 2016).

## Results

### Effects of Prior Protein Restriction on Preference for Solid Foods Differing in Protein Content

Data from the second 10-day period of maintenance diet exposure from Experiment 2 of Dibattista and Holder (1998) are re-plotted in Figure 1. Several important features can be observed. First, carbohydrate-rich test diet consumption was low in all groups (Figure 1A). Second, rats that had been previously protein-restricted but were currently fed a complete diet (PR-NR group) began the second phase eating more protein-rich test diet than rats that had never been protein-restricted (NR-NR; Figure 1B, compare red triangles to red diamonds) and those that were switched to the protein-free maintenance diet in the second phase (NR-PR; Figure 1B, compare red triangles to pink diamonds). Although the amount of protein-rich test diet consumed in the PR-NR group declined across days, it remained higher for the PR-NR group compared to the NR-NR group through Day 10 (Figure 1B, compare red triangles to red diamonds). Third, rats that had been previously protein-restricted and had been returned to protein-free maintenance diet (PR-PR) exhibited an immediate high level of protein-rich test diet consumption as opposed to a gradual progressive increase in protein-rich test diet as observed in the newly protein-restricted rats (Figure 1B, compare pink triangles to pink diamonds).

**Figure 1.**
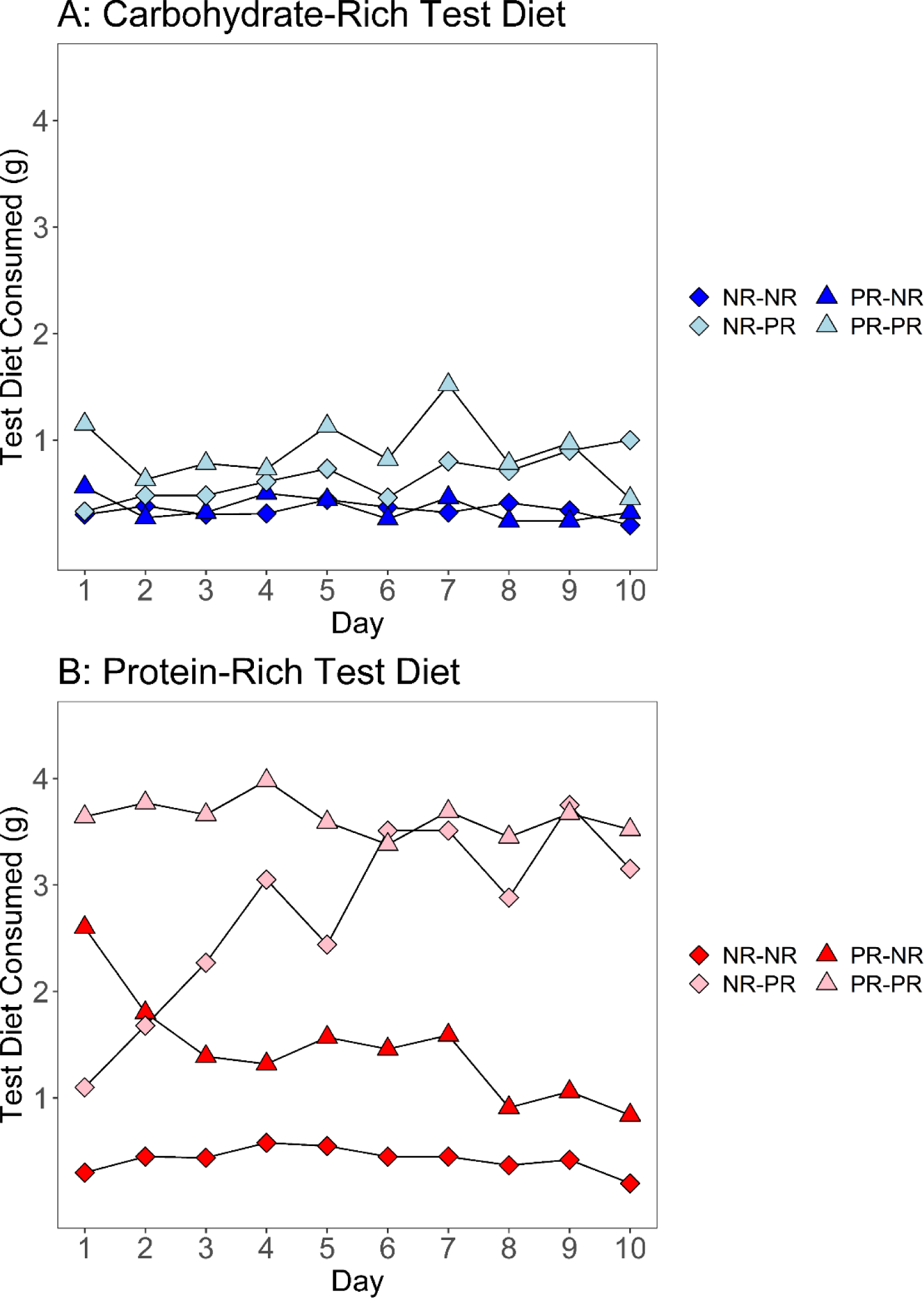
Re-analysis of data from Phase 2 of Experiment 2 of DiBattista and Holder (1998). Data were extracted from Figure 4 and re-plotted to compare amounts (g) of each test diet (**A**: consumption of carbohydrate-rich test diet = 94% dextrose; **B**: consumption of protein-rich test diet = 92% casein and 2% methionine) consumed between groups during the second phase of maintenance diet exposure. Maintenance diets were either protein-free (protein-restricted; PR) or nutritionally complete (not restricted; NR).

### Effects of Prior Protein Restriction on Preference for Solutions Differing in Protein Content

Figure 2A depicts the average grams of each solution consumed on each day of each preference assessment in the Torres et al. (2022) study. In the group of mice that started on the normal protein diet (Figure 2A, left panel, non-shaded area), mice consumed 4.6 – 7.1 grams of casein and 5.4 – 8.3 grams of maltodextrin solution across days of the first and second assessments, indicating a slight preference for maltodextrin. Following the switch to the low protein diet, the amounts of casein solution consumed increased dramatically to 11.1 – 13.2 g across days of the third and fourth assessments and the amounts of maltodextrin solution decreased slightly to 3.0 – 4.3 g across days of the third and fourth assessments (Figure 2A, left panel, shaded area). In the mice that started on a low protein diet, consumption of the casein solution substantially exceeded consumption of the maltodextrin solution by > 4-fold (Figure 2A, right panel, shaded area). After the switch to the normal protein diet, consumption of casein solution continued to exceed that of the maltodextrin solution for 28 days after protein normalization (Figure 2A, right panel, unshaded), although the difference was smaller than during the low protein diet period.

**Figure 2.**
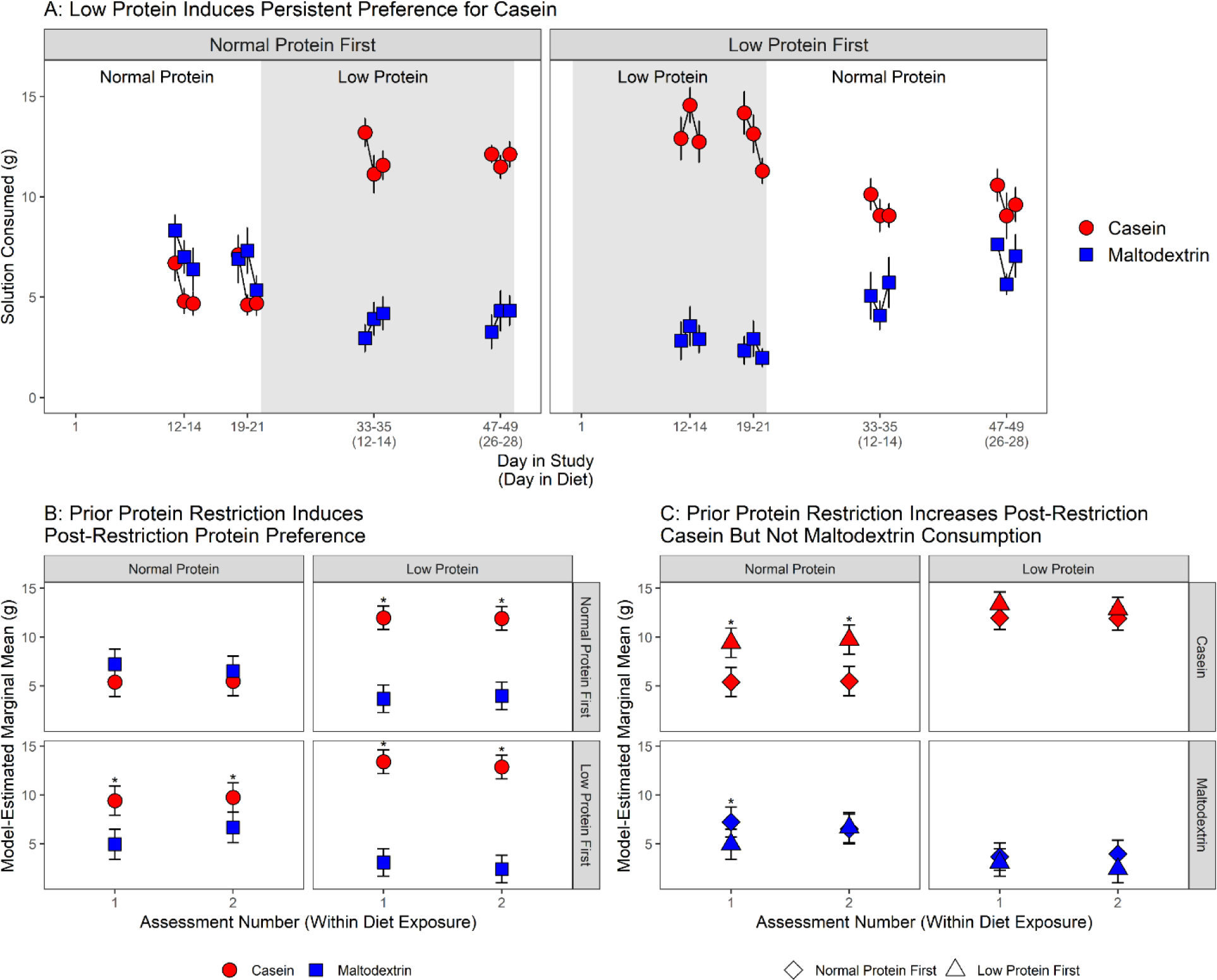
Re-analysis of data from Torres et al. (2022) cohort 2 preference assessments. **A**: Average daily amounts consumed of 4% casein and 4% maltodextrin solution in mice that were initially fed a diet consisting of 20% protein and subsequently a diet consisting of 5% protein (“Normal Protein First”) or the reverse (“Low Protein First”). **B**: Statistical comparisons between solutions of model-estimated marginal mean amounts consumed of each solution per assessment in Normal Protein First and Low Protein First mice during Normal Protein and Low Protein diet feeding. C: Statistical comparisons between groups of model-estimated marginal mean amounts consumed of each solution in Normal Protein First and Low Protein First mice during Normal Protein and Low Protein diet feeding. Each data point in **A** represents the average across mice. Each data point in **B** and **C** represents the model-estimated marginal mean for each combination of predictors. Error bars in **A** represent ±1 standard error of the mean (SEM). Error bars in **B** and **C** represent 95% confidence limits. * indicates a statistically significant (p < 0.05) difference between solutions (B) or groups (C).

After fitting a multilevel linear model to daily consumption amounts (complete model results presented in Table 1; estimated consumption amounts presented in Table 2), we used model-estimated marginal means to 1) compare amounts of casein consumed relative to amounts of maltodextrin consumed within each group (Figure 2B; Table 3); 2) compare amounts of casein consumed in the Low Protein First mice compared to amounts of casein consumed in the Normal Protein First mice (Figure 2C, top row; Table 3); and 3) compare amounts of maltodextrin consumed in the two groups of mice (Figure 2C, bottom row; Table 4). For the Normal Protein First group, mice consumed slightly more maltodextrin than casein during normal protein feeding (Figure 2B, top row, left) and much more casein than maltodextrin during low protein diet feeding (Figure 2B, top row, right). For the Low Protein First mice, amounts of casein consumed far exceeded amounts of maltodextrin consumed during low protein diet feeding (Figure 2B, bottom row, right). Following a switch to the normal protein diet, the difference between casein and maltodextrin consumption decreased but casein consumption remained higher than maltodextrin consumption (Figure 2B, bottom row, left). Finally, comparisons of the two groups of mice indicate that differences in consumption of casein versus maltodextrin during normal protein diet feeding were primarily driven by higher levels of casein consumption in the Low Protein First mice compared to the Normal Protein First mice and not by differences in maltodextrin consumption (Figure 2C, top left vs. bottom left).

**Table 1.** Regression estimates from linear multilevel model of daily amounts of casein and maltodextrin consumed. Predictor, estimate of the predictor (Estimate), standard error of the estimate (SE), resulting t-statistic (t), degrees of freedom (df; estimated via Satterthwaite method), and resulting p-value for the estimate (p; probability of the estimate under the null hypothesis that the true value of the estimate is 0). The fixed effects portion of the model included the predictors of Solution (Casein or Maltodextrin; dummy-coded with casein as the reference value), Diet (Low Protein or Normal Protein; dummy-coded with Normal Protein as the reference value), Group (Low Protein First or Normal Protein First; dummy-coded with Normal Protein First as the reference value), and Assessment No (continuous), and all possible interactions of the predictors. The random effects portion of the model allowed the coefficient of Solution and Diet to vary by mouse.

| Predictor | Estimate | SE | Statistic | df | p |
| --- | --- | --- | --- | --- | --- |
| Intercept | 5.3083 | 1.0523 | 5.0445 | 86.0 | < 0.0001 |
| Solution [Maltodextrin] | 2.6292 | 1.4337 | 1.8338 | 116.2 | 0.0692 |
| Diet [Low Protein] | 6.7167 | 1.2358 | 5.4349 | 326.3 | < 0.0001 |
| Group [Low Protein First] | 3.7792 | 1.4882 | 2.5395 | 86.0 | 0.0129 |
| Assessment No | 0.0833 | 0.5438 | 0.1532 | 325.0 | 0.8783 |
| Solution [Maltodextrin]:Diet [Low Protein] | -11.2542 | 1.7197 | -6.5442 | 325.0 | < 0.0001 |
| Solution [Maltodextrin]:Diet [Low Protein] | -8.4756 | 2.0294 | -4.1765 | 116.5 | < 0.0001 |
| Diet [Low Protein]:Group [Low Protein First] | -1.8708 | 1.7478 | -1.0704 | 326.3 | 0.2852 |
| Solution [Maltodextrin]:Assessment No | -0.7917 | 0.7691 | -1.0294 | 325.0 | 0.3041 |
| Diet [Low Protein]:Assessment No | -0.1417 | 0.7691 | -0.1842 | 325.0 | 0.8540 |
| Group [Low Protein First]:Assessment No | 0.2458 | 0.7691 | 0.3196 | 325.0 | 0.7494 |
| Solution [Maltodextrin]:Diet [Low Protein]:Group [Low Protein First] | 6.9464 | 2.4335 | 2.8545 | 325.0 | 0.0046 |
| Solution [Maltodextrin]:Diet [Low Protein]:Assessment No | 1.1333 | 1.0876 | 1.0420 | 325.0 | 0.2982 |
| Solution [Maltodextrin]:Group [Low Protein First]:Assessment No | 2.1797 | 1.0909 | 1.9982 | 325.2 | 0.0465 |
| Diet [Low Protein]:Group [Low Protein First]:Assessment No | -0.7208 | 1.0876 | -0.6628 | 325.0 | 0.5080 |
| Solution [Maltodextrin]:Diet [Low Protein]:Group [Low Protein First]:Assessment No | -2.6714 | 1.5404 | -1.7342 | 325.1 | 0.0838 |

**Table 2.** Model-estimated marginal mean amounts (g) of casein and maltodextrin consumed. Estimated mean amounts consumed per Solution, Diet, Group, and Assessment (Assessment No) were calculated based on the statistical model described in Table 1. Standard error of the estimate (SE) and lower and upper 95% confidence limits are also provided. using the emmeans command of the emmeans package. Post-hoc comparisons between groups, within each cohort, of intercepts (starting body weight) and slopes (change in body weight per day) within each phase (Treatment, Post-Treatment, and High-Fat Diet). Estimate of the model parameter for the first group in the comparison (“Est. 1”) and the second group in the comparison (“Est. 2”) along with the standard error (SE) of the estimate are provided for the intercept (starting body weight) and slope (coefficient of PND) parameters of the model. The emmeans command was used to conduct post-hoc, pairwise comparisons, and resulting t-statistic values [degrees of freedom] and associated p-values provided in the Comparison column for each model parameter.

| **Solution** | **Diet** | **Group** | **Assessment**<br>**No** | **Estimate** | **SE** | **Lower**<br>**95% CL** | **Upper**<br>**95% CL** |
| --- | --- | --- | --- | --- | --- | --- | --- |
| Casein | Normal Protein | Normal Protein First | 1 | 5.39 | 0.72 | 3.90 | 6.89 |
| Maltodextrin | Normal Protein | Normal Protein First | 1 | 7.23 | 0.74 | 5.69 | 8.77 |
| Casein | Low Protein | Normal Protein First | 1 | 11.97 | 0.59 | 10.75 | 13.18 |
| Maltodextrin | Low Protein | Normal Protein First | 1 | 3.68 | 0.68 | 2.28 | 5.09 |
| Casein | Normal Protein | Low Protein First | 1 | 9.42 | 0.72 | 7.92 | 10.91 |
| Maltodextrin | Normal Protein | Low Protein First | 1 | 4.96 | 0.74 | 3.42 | 6.50 |
| Casein | Low Protein | Low Protein First | 1 | 13.40 | 0.59 | 12.19 | 14.61 |
| Maltodextrin | Low Protein | Low Protein First | 1 | 3.10 | 0.68 | 1.69 | 4.50 |
| Casein | Normal Protein | Normal Protein First | 2 | 5.47 | 0.72 | 3.98 | 6.97 |
| Maltodextrin | Normal Protein | Normal Protein First | 2 | 6.52 | 0.74 | 4.98 | 8.06 |
| Casein | Low Protein | Normal Protein First | 2 | 11.91 | 0.59 | 10.70 | 13.12 |
| Maltodextrin | Low Protein | Normal Protein First | 2 | 3.97 | 0.68 | 2.56 | 5.37 |
| Casein | Normal Protein | Low Protein First | 2 | 9.75 | 0.72 | 8.25 | 11.24 |
| Maltodextrin | Normal Protein | Low Protein First | 2 | 6.68 | 0.74 | 5.13 | 8.22 |
| Casein | Low Protein | Low Protein First | 2 | 12.87 | 0.59 | 11.65 | 14.08 |
| Maltodextrin | Low Protein | Low Protein First | 2 | 2.41 | 0.68 | 1.01 | 3.82 |

**Table 3.** Comparisons of the amounts of each solution consumed (Casein – Maltodextrin). Comparisons between the amount of casein and maltodextrin consumed were conducted for each assessment (Assessment No) conducted within each Diet exposure for each Group. The difference in casein and maltodextrin consumed (Difference), standard error of the difference (SE), and degrees of freedom (df), t-statistic (t), and p value (p) for the difference are provided. Comparisons were conducted using the emmeans command of the emmeans package in the R programming language.

| Assessment No | Diet | Group | Difference | SE | df | t | p |
| --- | --- | --- | --- | --- | --- | --- | --- |
| 1 | Normal Protein | Normal Protein First | -1.84 | 0.93 | 25.03 | -1.97 | 0.0604 |
| 2 | Normal Protein | Normal Protein First | -1.05 | 0.93 | 25.03 | -1.12 | 0.2735 |
| 1 | Low Protein | Normal Protein First | 8.28 | 0.93 | 25.03 | 8.87 | < 0.0001 |
| 2 | Low Protein | Normal Protein First | 7.94 | 0.93 | 25.03 | 8.50 | < 0.0001 |
| 1 | Normal Protein | Low Protein First | 4.46 | 0.93 | 25.03 | 4.77 | < 0.0001 |
| 2 | Normal Protein | Low Protein First | 3.07 | 0.94 | 25.42 | 3.27 | 0.0031 |
| 1 | Low Protein | Low Protein First | 10.30 | 0.93 | 25.03 | 11.03 | < 0.0001 |
| 2 | Low Protein | Low Protein First | 10.45 | 0.93 | 25.03 | 11.19 | < 0.0001 |

**Table 4.** Comparisons between groups (Low Protein First – Normal Protein First) of the amounts of each solution consumed. Comparisons between groups in the amount of casein and maltodextrin consumed were conducted for each assessment (Assessment No) conducted within each Diet exposure for each Solution. The difference in amount consumed (Difference), standard error of the difference (SE), and degrees of freedom (df), t-statistic (t), and p value (p) for the difference are provided. Comparisons were conducted using the emmeans command of the emmeans package in the R programming language.

| Assessment No | Diet | Solution | Contrast | SE | df | t | p |
| --- | --- | --- | --- | --- | --- | --- | --- |
| 1 | Normal Protein | Casein | -4.03 | 1.02 | 20.73 | -3.96 | 0.0007 |
| 1 | Normal Protein | Maltodextrin | 2.27 | 1.04 | 20.24 | 2.18 | 0.0415 |
| 2 | Normal Protein | Casein | -4.27 | 1.02 | 20.73 | -4.20 | 0.0004 |
| 2 | Normal Protein | Maltodextrin | -0.15 | 1.05 | 20.49 | -0.15 | 0.8839 |
| 1 | Low Protein | Casein | -1.43 | 0.83 | 25.81 | -1.72 | 0.0976 |
| 1 | Low Protein | Maltodextrin | 0.59 | 0.96 | 21.89 | 0.61 | 0.5460 |
| 2 | Low Protein | Casein | -0.96 | 0.83 | 25.81 | -1.15 | 0.2610 |
| 2 | Low Protein | Maltodextrin | 1.55 | 0.96 | 21.89 | 1.62 | 0.1190 |

In the first preference assessment reported in the Naneix et al. (2020) study, rats initially fed the normal protein diet exhibited a preference for casein of ∼31% (i.e., they consumed more maltodextrin than casein) whereas rats initially fed the low protein diet exhibited a strong preference for casein of ∼75% (data are not replotted here but percentage preference values were extracted using PlotDigitizer on Figure 5 of Naneix et al). Following the diet switch, rats switched to low protein exhibited a progressive increase in casein preference, with preference at 64% on the first post-switch assessment and then 76% on the second, post-switch assessment. In comparison, rats switched to the normal protein diet (previously on low protein) exhibited a decline in casein preference to ∼67% initially and then to ∼55% at the second post-switch assessment. Thus, these data indicate that rats never exposed to low protein diet exhibit a ∼31% preference for casein, while rats previously exposed to low protein diet continued to exhibit a mild (∼55%) preference for casein up to 14 days after normalization of protein intake.

In the Chiacchierini et al. (2021) study, rats initially fed a normal protein diet (Normal Protein First) exhibited an ∼2-fold greater number of licks on the maltodextrin sipper than on the casein sipper (Figure 3A, top right, compare squares to circles in non-shaded area; complete model results presented in Table 5 and contrasts provided in Table 6). After the subsequent switch to a low protein diet, the NP First rats emitted more licks on the casein sipper compared to the maltodextrin sipper, with number of licks on the casein sipper tube increasing across the two assessments that followed the diet switch (Figure 3A, top right, compare circles to squares in shaded area). In comparison, rats initially fed a low protein diet (Low Protein First), exhibited a >4-fold greater number of licks on the casein sipper than the maltodextrin sipper (Figure 3A, top left, compare circle to square in shaded area). Following the switch to the normal protein diet, the number of licks on the casein sipper decreased and the number of licks on the maltodextrin sipper increased but casein licks remained greater than maltodextrin on the first assessment after the switch and licks on the two sippers were about equal on the last assessment (Figure 3A, top left, compare circles to squares in shaded area).

**Figure 3.**
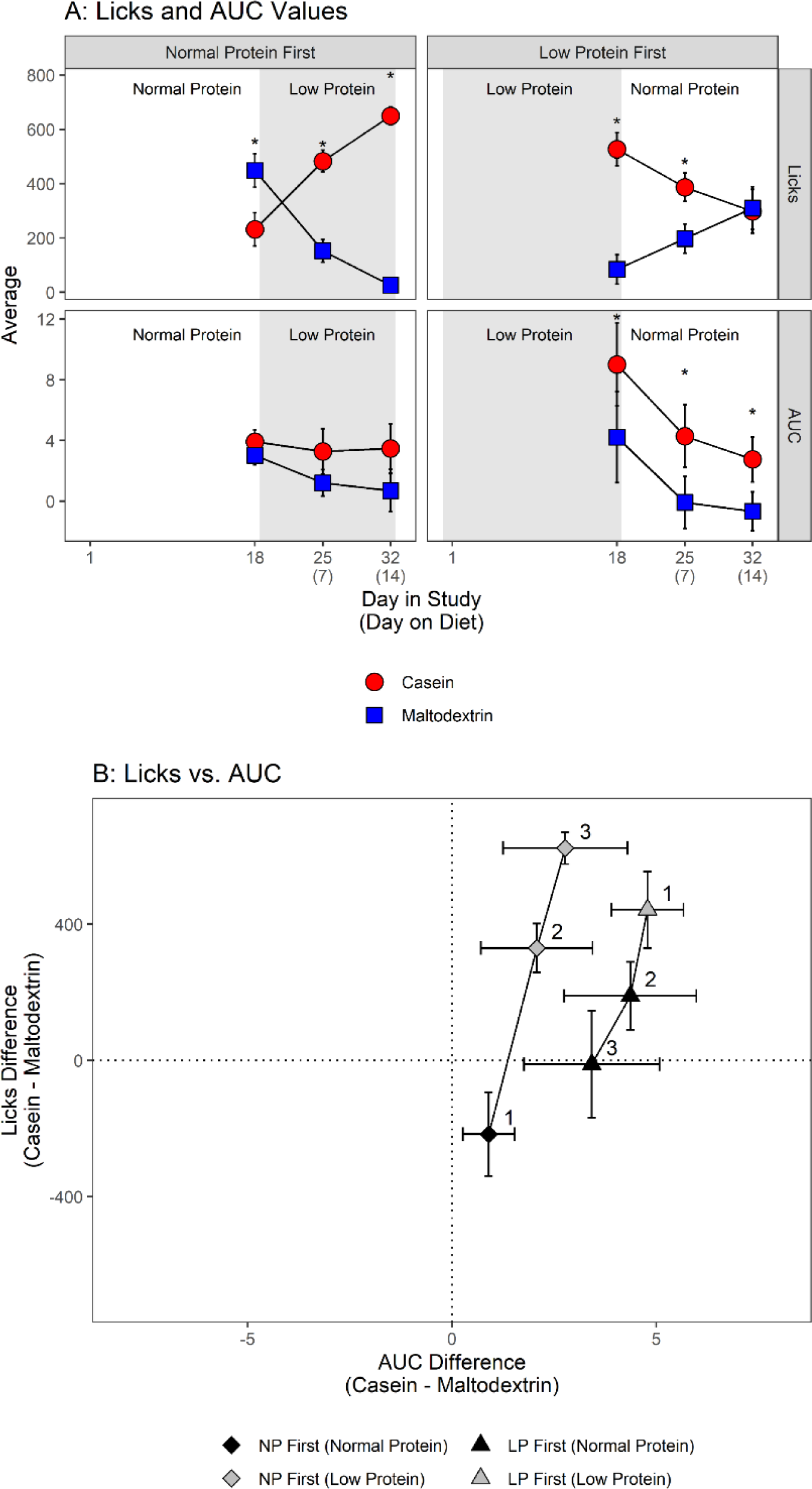
Re-analysis of data from Chiacchierini et al. (2021) preference assessments and VTA calcium imaging measured with fiber photometry. **A**: Number of licks on casein and maltodextrin sipper tubes (top row) in the group of rats that started on a normal protein diet (Normal Protein First) and rats that started on a normal protein diet (Low Protein First). Area-under-the-curve (AUC) values (bottom row) calculated from calcium influx signals in the VTA following licks on the casein or maltodextrin sipper tubes for each of the two groups of rats. **B**: Comparison of the association between the difference in licks for casein versus maltodextrin as a function of the difference in AUC values obtained following licks for casein versus maltodextrin. Each data point represents the average across mice. Numbers next to the data points indicate the assessment order within each group. Error bars represent ±1 standard error of the mean (SEM). * indicates a statistically significant (p < 0.05) comparisons between licks (A, top row) or AUC values (A, bottom row).

**Table 5.**
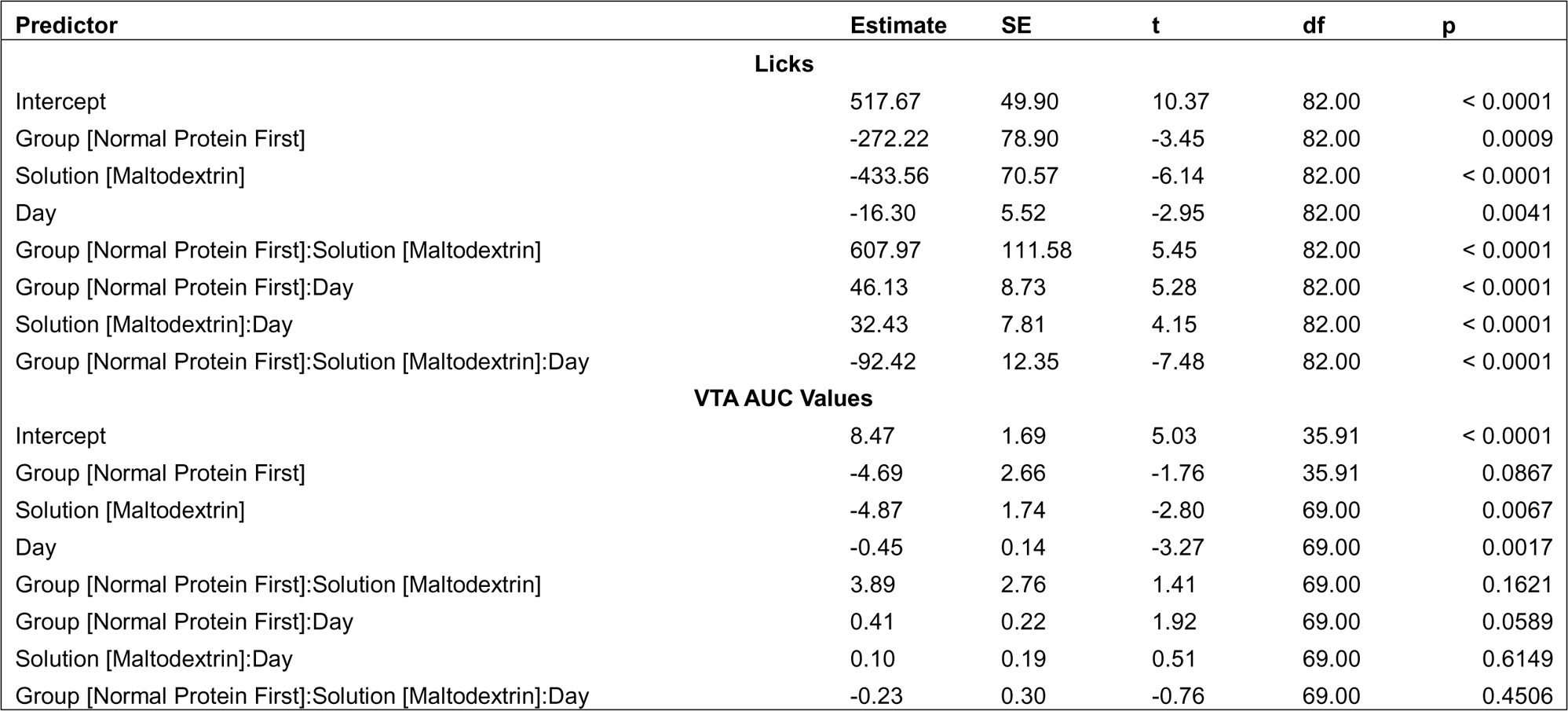
Regression estimates from linear multilevel model of licks for casein and maltodextrin and AUC values obtained from fiber photometry signals in VTA following licks on casein or maltodextrin sippers. Predictor, estimate of the predictor (Estimate), standard error of the estimate (SE), resulting t-statistic (t), degrees of freedom (df; estimated via Satterthwaite method), and resulting p-value for the estimate (p; probability of the estimate under the null hypothesis that the true value of the estimate is 0). The fixed effects portion of the model included the predictors of Solution (Casein or Maltodextrin; dummy-coded with casein as the reference value), Group (Low Protein First or Normal Protein First; dummy-coded with Low Protein First as the reference value), and Day (0, 7, 14; continuous), and all possible interactions of the predictors. The random effects portion of the model allowed the intercept to vary by mouse.

| Predictor | Estimate | SE | t | df | p |
| --- | --- | --- | --- | --- | --- |
| **Licks** |  |  |  |  |  |
| Intercept | 517.67 | 49.90 | 10.37 | 82.00 | < 0.0001 |
| Group [Normal Protein First] | -272.22 | 78.90 | -3.45 | 82.00 | 0.0009 |
| Solution [Maltodextrin] | -433.56 | 70.57 | -6.14 | 82.00 | < 0.0001 |
| Day | -16.30 | 5.52 | -2.95 | 82.00 | 0.0041 |
| Group [Normal Protein First]:Solution [Maltodextrin] | 607.97 | 111.58 | 5.45 | 82.00 | < 0.0001 |
| Group [Normal Protein First]:Day | 46.13 | 8.73 | 5.28 | 82.00 | < 0.0001 |
| Solution [Maltodextrin]:Day | 32.43 | 7.81 | 4.15 | 82.00 | < 0.0001 |
| Group [Normal Protein First]:Solution [Maltodextrin]:Day | -92.42 | 12.35 | -7.48 | 82.00 | < 0.0001 |
| **VTA AUC Values** |  |  |  |  |  |
| Intercept | 8.47 | 1.69 | 5.03 | 35.91 | < 0.0001 |
| Group [Normal Protein First] | -4.69 | 2.66 | -1.76 | 35.91 | 0.0867 |
| Solution [Maltodextrin] | -4.87 | 1.74 | -2.80 | 69.00 | 0.0067 |
| Day | -0.45 | 0.14 | -3.27 | 69.00 | 0.0017 |
| Group [Normal Protein First]:Solution [Maltodextrin] | 3.89 | 2.76 | 1.41 | 69.00 | 0.1621 |
| Group [Normal Protein First]:Day | 0.41 | 0.22 | 1.92 | 69.00 | 0.0589 |
| Solution [Maltodextrin]:Day | 0.10 | 0.19 | 0.51 | 69.00 | 0.6149 |
| Group [Normal Protein First]:Solution [Maltodextrin]:Day | -0.23 | 0.30 | -0.76 | 69.00 | 0.4506 |

**Table 6.** Comparisons between solutions of the AUC values from fiber photometry signals and the number of licks. Comparisons between solutions were conducted for each assessment, using the Day variable (0 for the first assessment, 7 for the second assessment, and 14 for the third assessment) and Group (Low Protein First or Normal Protein First). The difference in AUC values or licks (Difference), standard error of the difference (SE), and degrees of freedom (df), t-statistic (t), and p value (p) for the difference are provided. Comparisons were conducted using the emmeans command of the emmeans package in the R programming language.

| Group | Day | Difference | SE | df | t | p |
| --- | --- | --- | --- | --- | --- | --- |
| **Licks (Casein - Maltodextrin)** |  |  |  |  |  |  |
| Low Protein First | 0 | 433.56 | 70.57 | 69.00 | 6.14 | < 0.0001 |
| Normal Protein First | 0 | -174.42 | 86.43 | 69.00 | -2.02 | 0.0475 |
| Low Protein First | 7 | 206.56 | 44.63 | 69.00 | 4.63 | < 0.0001 |
| Normal Protein First | 7 | 245.50 | 54.66 | 69.00 | 4.49 | < 0.0001 |
| Low Protein First | 14 | -20.44 | 70.57 | 69.00 | -0.29 | 0.7729 |
| Normal Protein First | 14 | 665.42 | 86.43 | 69.00 | 7.70 | < 0.0001 |
| **AUC (Casein - Maltodextrin)** |  |  |  |  |  |  |
| Low Protein First | 0 | 4.87 | 1.74 | 69.00 | 2.80 | 0.0067 |
| Normal Protein First | 0 | 0.98 | 2.13 | 69.00 | 0.46 | 0.6475 |
| Low Protein First | 7 | 4.19 | 1.10 | 69.00 | 3.80 | 0.0003 |
| Normal Protein First | 7 | 1.92 | 1.35 | 69.00 | 1.42 | 0.1600 |
| Low Protein First | 14 | 3.51 | 1.74 | 69.00 | 2.01 | 0.0480 |
| Normal Protein First | 14 | 2.85 | 2.13 | 69.00 | 1.34 | 0.1855 |

### Physiology

Chiacchierini et al. (2021) used fiber photometry to measure calcium influx in ventral tegmental area (VTA) neurons when rats licked the protein and maltodextrin sipper tubes. They found that in rats initially exposed to a normal protein diet and then switched to a low protein diet, the AUC values obtained during normal protein diet were similar (Figure 2A, bottom left, compare circles to squares in non-shaded area). Following the switch to the low protein diet, AUC values were slightly higher for casein, primarily because the values for maltodextrin declined (Figure 2A, bottom left, compare circles to squares in shaded area). In rats initially fed a low protein diet (Low Protein First), the area-under-the-curve (AUC) values calculated from the photometry signals obtained immediately following licks on the casein bottle were higher, on average, than those immediately following licks on the maltodextrin bottle (Figure 3A, bottom right, compare circle to square, shaded area; complete model results provided in Table 5 and contrasts in Table 6). Following the diet switch, rats that were previously fed the low protein diet continued to exhibit higher average AUC values for casein than maltodextrin (Figure 3A, bottom right, compare circles and squares in non-shaded area).

Finally, for each rat, we calculated the difference in licks for casein vs. maltodextrin and the difference in AUC values for casein vs. maltodextrin for each preference assessment. We then calculated the average difference in licks and average difference in AUC values. Figure 3B compares the association between the difference in licks and the difference in AUC values for the Normal Protein First and Low Protein First rats. As the difference in AUC value increased, the difference in licks increased in each group, but the results for the Low Protein First group are shifted rightward compared to the Normal Protein First group (Figure 3B, compare triangles to diamonds).

## Discussion

Re-analysis of our own and others’ data on preference for protein versus carbohydrate during periods of exposure to low and normal protein diets indicates that dietary protein restriction induces persistent increases in relative and absolute consumption of protein. Further, re-analysis of other data indicates that dietary protein restriction also exerts a persistent effect on VTA activity in response to protein versus carbohydrate. Finally, our re-analysis indicates that prior protein restriction shifts rightward the relation between relative behavior (licks) and relative VTA activity with respect to protein and carbohydrate. Findings that dietary protein restriction exerts effects on preference and brain activity that persist for an extended period following normalization of protein intake suggests that relative and absolute protein consumption is driven jointly by prior experience and current protein need and carries important implications for our understanding of protein motivation and how we investigate the determinants of protein motivation.

That prior dietary protein restriction may exert persistent influences on behavior and physiology raises several experimental questions. From a behavioral standpoint, we know little about the conditions under which persistent effects occur. Obvious questions of interest are whether induction of persistent effects depends on the duration or degree of protein restriction. Further, although the present analyses indicate that prior protein restriction can exert an influence on protein preference for up to 4 weeks (see Figure 2A), it is unknown whether extended exposure to normal protein intake would eventually result in relative and absolute consumption levels of protein versus carbohydrate observed in animals that have never been protein restricted. Additionally, some studies (Chiacchierini et al., 2021; Naneix et al., 2020) utilize individual conditioning sessions in which the animals receive access to only one solution prior to preference assessments and the impact of such exposure to individual macronutrients on subsequent preference following prior protein restriction remains to be investigated. Finally, from a protein leverage standpoint these results suggest that the intake target for protein may be increased by prior protein restriction, at least for a period of weeks, which could be investigated within the nutritional geometry framework (Raubenheimer & Simpson, 2019).

From a biological standpoint, we know little about the underlying changes that mediate persistent effects of prior protein restriction. A substantial body of data indicates that the behavioral and physiological changes that occur during dietary protein restriction depend on actions of liver-derived FGF21 in the brain (for a review see Hill et al., 2020). It is assumed, but to our knowledge has not been demonstrated, that FGF21 levels decrease once protein intake normalizes. If so, then it may be that FGF21 induces downstream adaptations that underlie the persistent effects of prior protein restriction. As an example, it may be that FGF21 induces long-lasting or permanent changes in brain responses to protein or protein-associated cues that underlie the influence of prior protein restriction. Or, perhaps, liver-derived production of FGF21 persists for an extended period following prior protein restriction and these high levels of FGF21 mediate the persistent effects of prior protein restriction. Finally, the finding that some effects of dietary protein restriction, such as changes in the rate of body weight gain adjust rapidly and appear to completely reverse (Torres et al. 2020), whereas other effects, such as preference, appear to be sensitive to prior conditions, suggests there must be distinct mechanisms, that remain to be discovered, underlying those different outcomes.

In conclusion, persistent effects of prior dietary protein restriction indicate that behavior and possibly physiological responses to protein depend on prior dietary conditions to an extent previously not recognized, in addition to current dietary conditions. Experiments that vary dietary protein intake within animals can determine how prior and current nutritional conditions interact to govern relative and absolute intake of protein and other macronutrients as well as what physiological changes mediate the effects of current versus prior nutritional conditions (for a related discussion of dealing with irreversible effects, see Sidman, 1960, pp. 52-53). Investigation of these questions will be able to leverage modern genetic and neurological tools to disentangle the mechanisms that mediate persistent responses from initial responses to dietary protein restriction. Finally, to the degree that beneficial effects of dietary protein restriction can be extended beyond the period of dietary protein restriction, findings may be leveraged to the benefit of human health.

